# Social genetic effects on feed intake in pigs: from feeding visits to total intake

**DOI:** 10.64898/2026.09.28.755174

**Authors:** P. Nuñez, Z.G. Vitezica, A. Legarra, A. Zubiri-Gaitán, S. Negro Rama, J. Casellas, L. Varona, N. Ibáñez-Escriche

## Abstract

Social genetic effects can substantially influence phenotypes in animals and plants and the study of them is of interest. Group housing in animals – for example, in pigs – provide an ideal framework for studying interactions among individuals. In animals, the genetic of social effects has been studied by simplifying the underlying interactions – for example, by assuming that all the individuals in the group contribute equally to the social environment. This homogeneous behaviour is not observed in real conditions. Here, we address this issue showing that identification of interacting individuals is feasible in pig behaviour during growth, and we propose a model that leads to more accurate modelling of social genetics. Our approach infers the strength of interaction via time interval between visits of animals to automatic feeders. This measure of strength was then used in a reaction norm model to evaluate the genetic determinism of social effects. We also build a framework to transform variance components estimated for a trait continuously measured, such as feed intake per visit, into variance of total or aggregated trait e.g. during the growth period. Including social effects improves the fitting of the models. Visit-based social interaction was highest at short time intervals, decreasing rapidly and becoming negligible after 80 seconds. Variance components for the aggregated trait – the total feed intake – support the contribution of the social interactions. Our approach allows more precise understanding of social effects through individual feeding events and a measure of interaction strength, and further translates these into in an aggregated phenotype of interest.

**Article summary:** Animals living in groups constantly interact with one another, and these interactions can influence their performance and behaviour. Conventional approaches often assume that all animals contribute equally to the social environment; however, it is not true in real conditions. In this study, we identified and measured interactions between pigs based on the time interval between their visits to automatic feeders. We incorporated this information into a genetic model to account for differences in the strength of social interactions. Our results show that social interactions are strongest when animals visit the feeder close together and can meaningfully affect overall feed intake during growth.

## Introduction

Groups of animals or plants interact among them. Each individual has a genotype that determines its own phenotype and, at the same time, may carry genes that influence the phenotypes of the individuals with whom it interacts within the group. The genetics of social interactions, known as social (or indirect) genetic effects, are not yet fully understood, despite a growing interest (Muir, 2005; Bijma et al., 2007; Baud et al., 2017; Salomon et al., 2026). Pigs offer an important case of study for social effects in livestock, as they are raised in “pens” where they exhibit social interactions, including competition for food. In these pens, social genetic effects of the pen mates may substantially influence growth and performance. For animal husbandry, ignoring these indirect effects may therefore lead to suboptimal responses to selection (Muir, 2005). Bijma et al. (2007) proposed a social genetic model in which the total breeding value is composed of both direct and indirect or social genetic effects. This framework has been successfully applied to several productive and behavioural traits in pigs (Bergsma et al., 2008a; Nielsen et al., 2018; Herrera-Cáceres et al., 2019; Leite et al., 2023). However, Bijma et al. (2007) assumes equal contributions of all contemporaries in the pen to the social environment, which may oversimplify the underlying interactions. However, pairs of interacting individuals can sometimes be identified; the most obvious case is maternal effects at growth in mammals (Willham, 1972) or Bijma’s hypothetical example of neighbouring trees (Bijma, 2011). In this work we show that identification of interacting individuals is feasible in pig behaviour during growth, and this leads to better modelling of social genetics.

Automatic feeding systems facilitate the identification of social interactions in pigs by providing detailed longitudinal data on feed intake and feeding behaviour. Automatic feeders record which animals enter the feeder, when do they enter, how much they eat, and when do they leave. Based on these data, previous studies proposed approaches that model social genetic effects using information from individual feeder visits (Angarita et al., 2021; Nuñez et al., 2025). In these models, each visit to the automatic feeder is modelled individually. The phenotype of the (so called “focal”) pig entering the feeder is affected by a social effect which is assumed to originate only from the pig entering the feeder immediately afterwards (hereinafter the “follower”). This approach is similar to a maternal-effect model (Willham, 1963; Cheverud, 1984), so the genetic social effect of the follower acts as an environmental effect on the focal animal at the feeder. Unlike Bijma et al. (2007) classical social genetic models, this framework focal-follower accounts for the frequency of interactions among animals across multiple visits and combinations of individuals. The intensity of the interaction is assumed to depend on the time interval between visits. A limitation of visit-based models proposed by Angarita et al. (2021) is that the trait analysed corresponds to individual feeding events, rather than to the economically relevant aggregated phenotype, such as feed intake over the entire testing period.

The objectives of the present study are two. First, we propose a visit-based approach for feed intake that incorporates time interval between feeder visits as a proxy for the strength of the interaction. This is modelled using random regression on a continuous measure of strength in a way similar to norm reaction models. Second, genetic end environmental variances and covariances are inferred from the individual visits, and then analytically aggregated to obtain global genetic variance components for the phenotype across the whole period, in this case, total feed intake. In this manner, we build a framework to model frequent interactions across possibly many pairs of animals, but resulting in a single phenotype of interest, which in this work is feed intake during a period but it could be any trait of interest that is continuously measured.

## Materials and methods

### Ethical review

All farm management practices were approved by the Ethics and Animal Welfare Committee of the Polytechnic University of Valencia, in accordance with Council Directives 98/58/EC and 2010/63/EU (reference number 2017/VSC/PEA/00212).

### Experimental data

The database used in this study belonged to a selection nucleus farm of Iberian pigs (IngaFood S.A., Almendralejo, Extremadura, Spain). Pigs were fed using automatic feeders (Nedap, Groenlo, the Netherlands). Pigs were housed in pens with an average of 12.8 pigs per pen with a single feeder per pen. Only visits to the feeder up to 180 days of age were considered. Animals entered the testing period with an average initial weight of 31.4 ± 8.4 kg and finished with 80.4 ± 12.0 kg, with an average testing duration of 85.5 ± 15.1 d. Automatic feeder records contained 1,776,299 visits from 1357 animals, recorded from August 2021 to September 2025. Each visit included corresponding animal identification (ID), feed intake and time of entrance and of leaving the feeder, from which duration per visit was obtained. Hereinafter, we call a pig visiting the feeder a “focal” pig, and the next pig entering, the “follower”. The original database was processed to include two additional variables for each visit: (i) the ID of the follower pig, and (ii) the time interval between the exit of the focal pig and the entry of the follower. Basic filtering was applied to remove abnormalities such as lack of ID, duplicated visits and outliers in feed intake or duration per visit. Visits were discarded when feed intake < 1g and duration < 2 s. Outliers in feed intake and visit duration were identified using an age-based slicing window approach. For each visit, a window of ±5 days around the animal’s age was defined. Within each window, the 0.99 percentile of feed intake and visit duration was calculated and values exceeding this percentile were removed. This approach allowed for age-specific outlier detection while preserving biologically meaningful variation. Pedigree information included 2,304 individual-sire-dam entries, and the median of the genetic relationship between pigs was 0.08.

### Statistical model

At the visit level, the model assumes that the direct additive genetic effect corresponds to the “focal” pig occupying the feeder, whereas the social effect corresponds to the follower pig (Angarita et al., 2021). Furthermore, we assume that the magnitude of this social effect decreases with a function *Φ* of the time interval between entry of the follower and the exit of its predecessor. To model feed intake at the visit level (FIV), we use a random regression model. We compared three different models, including different combinations of pen effects, direct genetic effects, social genetic effects, and permanent environmental effects as random factors, as follows:

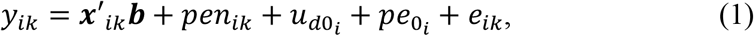

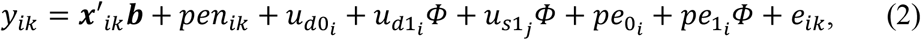

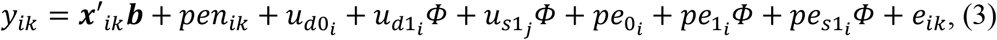

where *y*_*ik*_ is the feed intake of *i*^*th*^ individual on its *k*^*th*^ visit, ***x***_*ik*_ is the vector of fixed effects including the general mean, the hour of the day as a categorical variable, the age of the animal *i* at visit *k* as a covariate, and the covariate *Φ* as a gradient of social competition (see the definition below), *pen* is the random pen effect, *u*_*d*_ is the random direct additive genetic effect of the *i*^*th*^ focal pig, *u*_*s*_ is the random social additive genetic effect belonging to the *j*^*th*^ follower pig, and *pe* is the random permanent environmental effect of animal *i*. The direct additive effect belongs to the pig in the feeder, while the social effect belongs to the follower pig. Note that *u*_*d*_, and *pe* involve an intercept (*u*_*d*0_, and *pe*_0_), and *u*_*d*_, *u*_*s*_, and *pe* involve a slope (*u*_*d*1_, *u*_*s*1_ and *pe*_1_) which depends on the covariate *Φ*, which is a function of the time interval between the exit of the animal in the feeder and the entering of the following. Note that *u*_*s*_ does not include an intercept because the model assumes that the feed consumption of one animal at the feeder station only depends on the interval time in which the follower enters, being zero when the interval is too long.

These models can be interpreted from different perspectives. First, at the level of each visit, the phenotype of a given feeding event only depends on the animal consuming feed during that visit (Model 1). In Model 2, the phenotype also depends on the social genetic effect of the animal that immediately replaces it at the feeder. The social effect is mitigated by a covariate *Φ* in a manner similar to a norm reaction model. In Model 3 the permanent environmental effect of the follower is also considered.

From the perspective of an individual animal *i*, models 2 and 3 yield both a direct genetic value and a social genetic value for each animal. The direct genetic value is estimated from the visits in which animal *i* is the focal individual consuming feed: a slope of zero (*u*_*d*1_ = 0) indicates that the focal animal’s feed intake is unaffected by the social environment, whereas a slope (*u*_*d*1_) higher or lower than zero indicates that the animal consumes more or less feed depending on the social pressure exerted by its followers. The social genetic value, in contrast, is estimated from the feed intake of the focal (preceding) animals for whom animal *i* acts as the follower: a slope of zero (*u*_*s*1_ = 0) indicates that animal *i* does not affect the previous animal, whereas a slope (*u*_*s*1_) higher or lower than zero indicates that the animal preceding *i* is affected positively or negatively respectively. The random regression analysis also provides estimates of variance components, and breeding values at different points along the covariate *Φ*, which is a function of the time between “focal” and “follower” visits.

The following normal distributions are assumed for random effects. The pen effect was assumed to follow 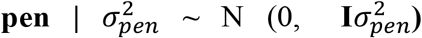, the additive genetic effects [***u***_***d*0**_ ***u***_***d*1**_ ***u***_***s*1**_]^′^| **G**_**0**_ ~ MVN (0, **G**_**0**_ ⊗ **A**), the permanent environmental effects [***pe***_**0**_ ***pe***_**1**_]^′^ | **P**_**0**_ ~ MVN (0, **P**_**0**_ ⊗ **I**) and the residuals 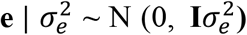, where **A** is the relationship matrix, **I** is an identity matrix with the same order of the number of records, 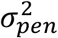 and 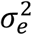 are the pen and residual effects variances, respectively, ⊗ denotes the Kronecker product, and **G**_**0**_ and **P**_**0**_ are the (co)variance matrices of the genetic and permanent environmental effects, respectively, as follows:

Model 2:

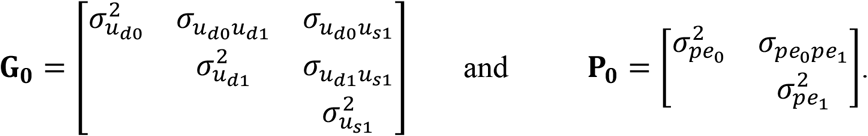

Model 3:

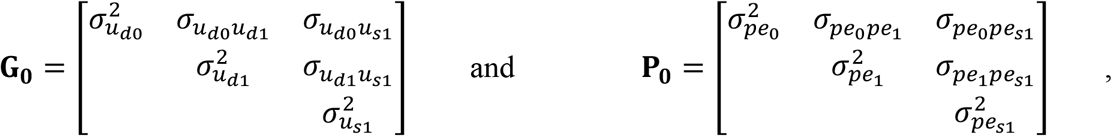

where *pe*_0_ and *pe*_1_ correspond to the intercept and slope of permanent environmental effect of the focal animal, while *pe*_*s*1_ refers to the slope of the permanent environmental effect of the follower.

### Definition of the covariate Φ

To describe the intensity of social interaction during feeder visits, the time interval between consecutive visits was transformed into a covariate (*Φ*) ranging from 0 to 1. This transformation was motivated by the assumption that social competition at the feeder does not vary linearly with time. Very short intervals between consecutive visits are expected to reflect strong social pressure, whereas beyond a certain threshold the effect of longer intervals should become biologically non-relevant. To define an appropriate function, the dataset was first divided into 8 time-interval classes (e.g. 0 s, 0-20 s, 10-30 s, 20-40s, 30-50s, 40-60s, 60-120s, and more than 120s). Preliminary estimates of social variance for each of the 8 intervals were obtained applying a simple model 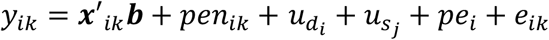 (note that this model does not include random regression analysis). Based on these empirical trends, a smooth function was fitted to approximate how social variance changes with increasing time interval between visits. This function was fitted not to the social variance itself, but to its square root, evaluated at standardized time interval points. This choice ensures consistency with the random regression model, in which variance components scale with the square of the covariate, i.e., *Var* = *f*(*Φ*)^2^. This fit resulted in a function defining covariate *Φ* as

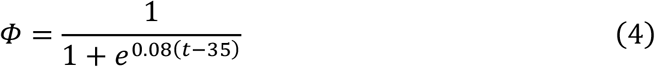

where *t* is the time interval (in seconds) between consecutive visits. Under this formulation, *Φ* approaches 1 for very short intervals (high social interaction) and tends to 0 for long intervals (>60 s), corresponding to negligible social interaction. Results from preliminary analysis of subdatasets and construction of covariate *Φ* is included in Supplementary material. Figure S1 illustrates variance component estimations on different subsets, while Figure S2 shows how the defined covariate *Φ* aligns with the square root of the normalized social variance.

### Variance component estimation

Variance and covariance components were estimated using restricted maximum likelihood (REML) with the Average Information algorithm implemented in the BLUPF90 software suite (Aguilar et al., 2018).

### Variance components over time intervals

Random regression analysis allows to assess how variance components (as well as individual breeding values) change over a gradient of social interaction (the covariate range). Once genetic (co)variances (**G**_**0**_) were obtained, additive variance at specific points of the covariate *Φ* can be obtained as 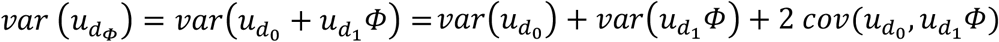, which is equivalent to 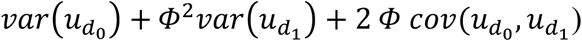. Social variance at one specific point can be obtained as 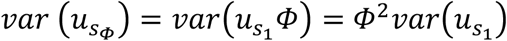. Permanent environmental variance can also be obtained at different points of the gradient, according to each model. Genetic direct-social covariance at point *Φ* was obtained as 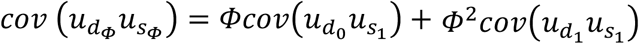. The proportion of the phenotypic variance explained by direct additive variance 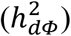 and social additive variance 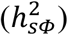 at specific point was calculated as 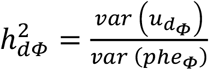 and 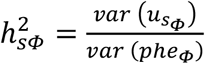, being 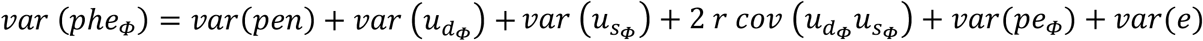. The term *r* is the mean relatedness coefficient between animals in the same pen, which is 0.08. This is because additive effects and social effects come from two different, but related, animals. Under this visit approach, additive direct breeding values 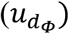 and social additive breeding value 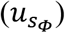 for animal *i* at point *Φ* are 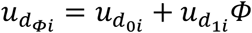 and 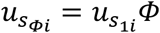, respectively. The total breeding value can also be obtained as 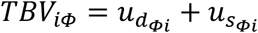. Note that this approach differs from the classic social approach (Bijma et al., 2007) where 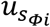 is multiplied by (*n* − 1) i.e. all possible followers. The difference comes because in the visit-based approach (this study), the social effect of each specific visit is assumed to come only from the follower, and not from all the (*n* − 1) pen mates.

### Genetic correlations over different time-intervals

To evaluate the impact of the time interval on rankings based on direct additive breeding values, we estimated the pairwise genetic correlation between pairs of different time intervals (0, 10, 20, 30, 60, 90, and 120) as follows:

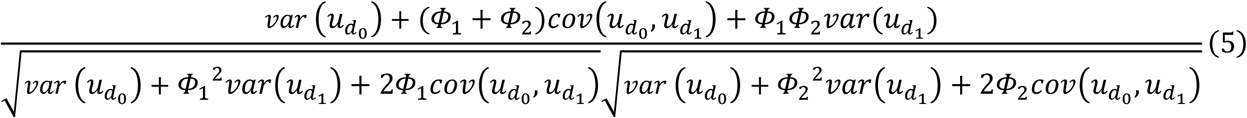

where *Φ*_1_ and *Φ*_2_ refer to the value of the covariate at different time points. The social genetic correlation between different time intervals was estimated as follows:

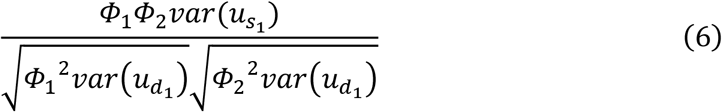

This analysis provides an indication of how similar the ranking would be if selection were based on breeding values estimated at one time interval compared with another.

### Variance component conversion into a total feed intake along the period

The visit-based approach allows a more precise characterization of social effects by explicitly assigning a single social interaction to each feeding event. However, a key limitation of the visit-based approach is that the resulting solutions (i.e., variance components and breeding values) are defined at the level of Feed Intake per Visit (FIV), which is not itself an economically relevant trait in pig production systems whereas the Total Feed Intake during the Period (TFIP) is of economic relevance, and a better descriptor of the biology. To address this limitation, we performed a transformation of the variance components estimated under the visit-based approach for FIV to approximate those corresponding to TFIP, in which a single phenotype is obtained as the sum of individual feed intakes across visits (i.e. TFIP is the sum of all FIV). The additive and social genetic variances are transformed from the FIV to the TFIP scale as follows:

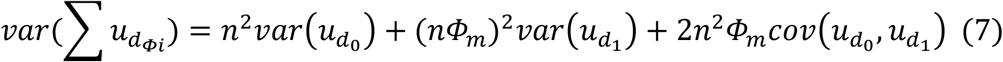

and

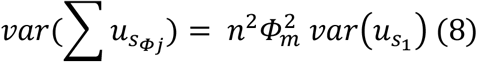

Where *n* is the median number of visits, and *Φ*_*m*_ the median value of the covariate *Φ* across the total visits. These assumptions aim to approximate the reconstructed variance of a “representative” animal (see the derivation of Eq. 7 and Eq. 8 in the Appendix). The same approach is used to recover permanent effects and pen effects in the TFIP scale. In the case of residual variance, it is reconverted as *var*(∑ *e*_*i*_) = ∑ *var*(*e*_*i*_) = *n var*(*e*_*i*_) because *cov*(*e*_*ik*,_*e*_*il*_) = 0. The total direct-social covariance was obtained as 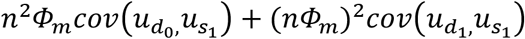.

## Results

### Variance component estimation of individual visits with social effect of the follower

Variance component estimates for the three models are presented in Table 1. Likelihood ratio tests (LRT) indicated that each increase in model complexity significantly improved model fit (Model 1 vs. Model 2: LRT = 57,630, *P* < 0.001; Model 2 vs. Model 3: LRT = 298, *P* < 0.001). When the permanent environmental effect of the follower was included (Model 3), a substantial proportion of the variance was captured by the follower permanent environmental effect and the estimated social genetic variance was markedly reduced. However, the asymptotic correlation obtained from the inverse Average Information matrix was −0.60 for the pair of variance components involving the social genetic and social permanent environmental effects, indicating a considerable degree of confounding genetic and environmental components in the likelihood. For this reason, hereinafter we describe mainly results from model 2, with the caveat that the social genetic variance may be overestimated, as it likely contains both genetic and environmental effects.

**Table 1.**
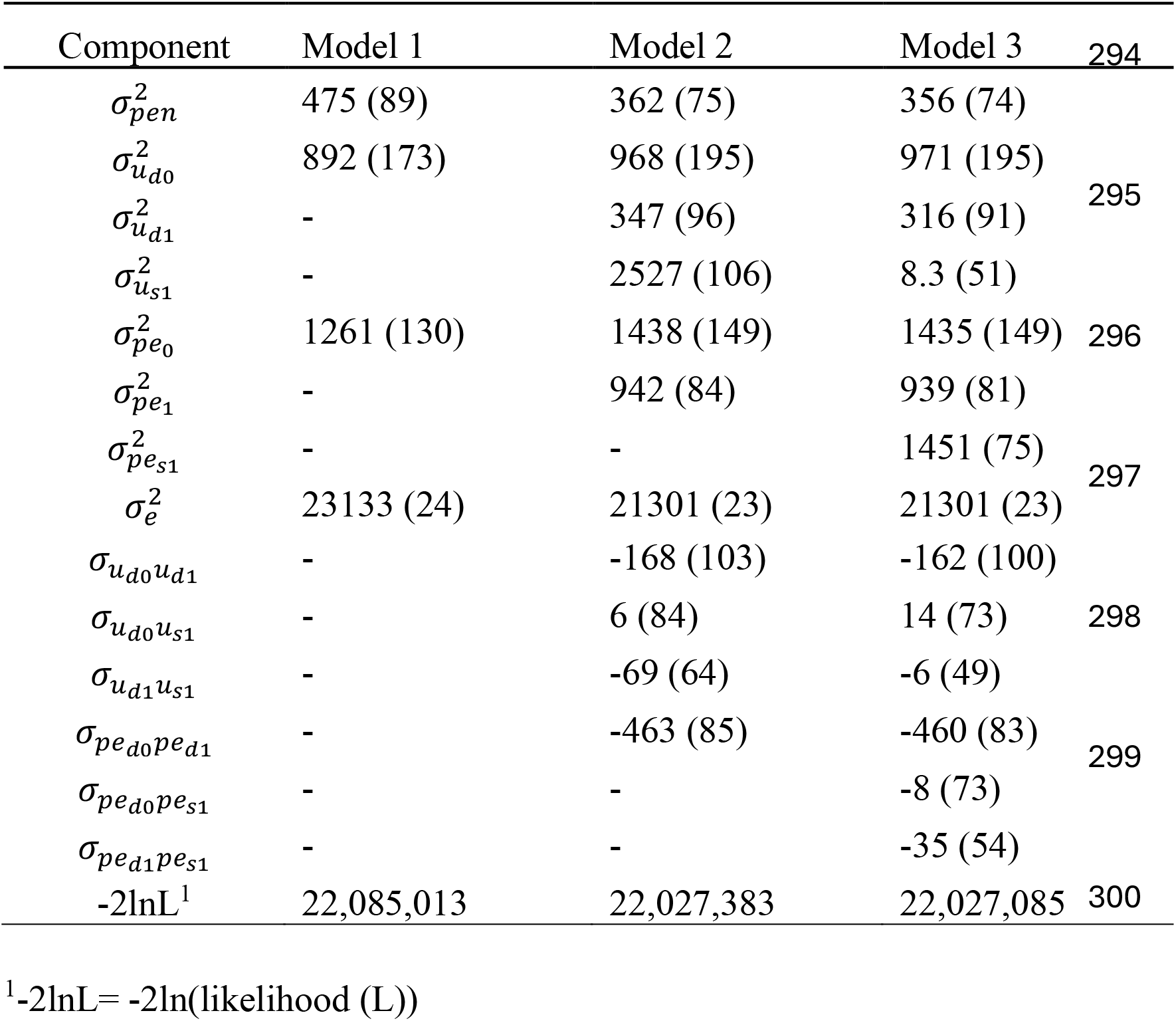
Variance component estimation for feed intake per visit (in grams) with their standard deviation (in brackets) using the three proposed models.

Figure 1 shows the proportion of the phenotypic variance explained by the direct additive genetic 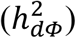, social additive genetic 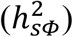, and permanent environmental (*pe*^2^) effects under Model 2 over the gradient due to time interval between focal and follower pigs. Social genetic variance accounted for up to 8.5% of the phenotypic variance when the time interval was zero, but declined rapidly and became negligible at approximately 60 s. In contrast, direct heritability remained relatively constant at around 3.5% across the entire gradient, whereas the permanent environmental effect explained approximately 5.3% of the phenotypic variance throughout. The correlation between direct and social additive genetic effects remained close to zero, ranging from −0.04 to 0.

**Figure 1.**
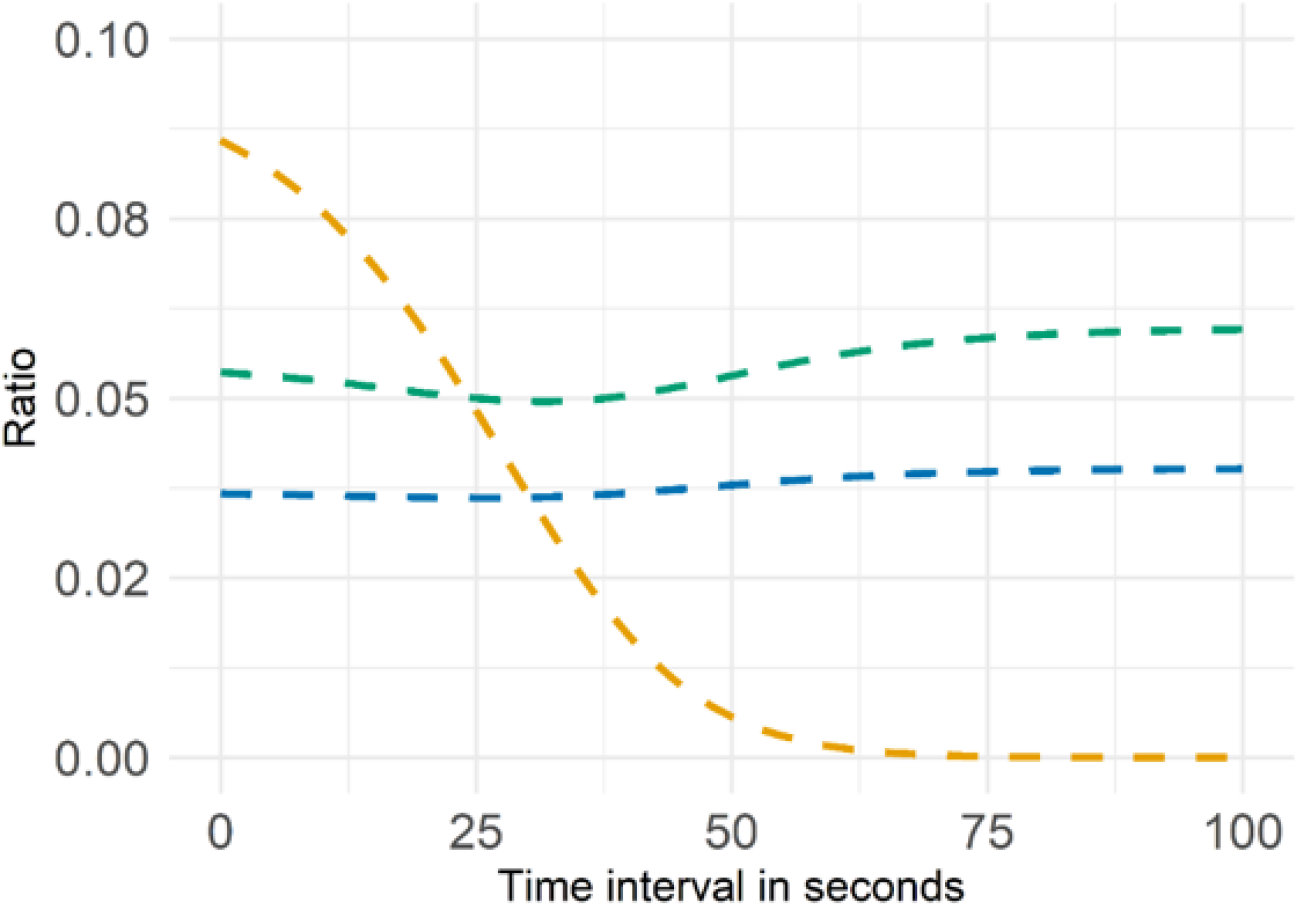
Proportion of the phenotypic variance explained by additive direct effect (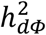, in blue), social genetic effect (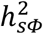, in yellow) and permanent effect of the focal animal (*pe*^2^, in green) over time interval.

### Correlations within direct genetic and total genetic effects between different time intervals

Direct additive genetic and total additive genetic correlations were assessed at different time interval lengths. For clarity, we describe only correlations between the first point (time 0) with the rest of the points, but all the correlations are shown in Figure 2. For additive genetic correlations, results denote high consistency between time points, with a range between 1 and 0.92. On the other hand, total additive correlations (including both additive and social components) showed a range between 1 and 0.62, presenting an abrupt reduction between 30 and 60 s (from 0.98 to 0.74). Social correlations are not shown. This is because models 2 and 3 include only the slope but not an intercept for social effect. Therefore, pairwise social correlations between points are 1.

**Figure 2.**
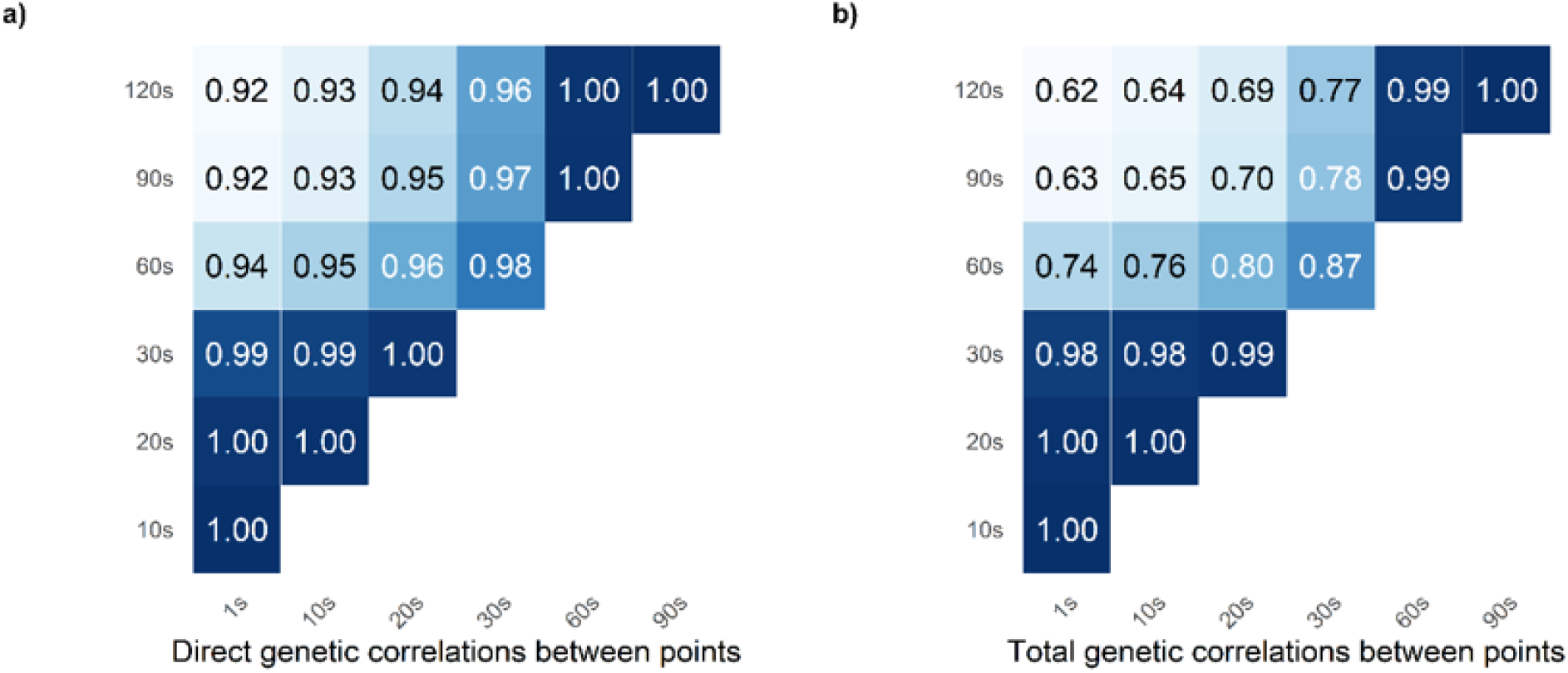
Direct additive genetic correlations (left) and total additive genetic correlations (right) between different time intervals (in seconds).

Figure 3 illustrates intra-animal variation on direct breeding values 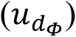 and social breeding values 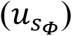 for 20 random animals. For 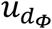, greater variation between animals was observed after 25 s, whereas for 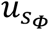 the differences between animals were markedly larger at the beginning. Note that 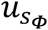 converges to 0 for every animal when the interval time approximates 90 seconds, because the value of the *Φ* function is nearly 0.

**Figure 3.**
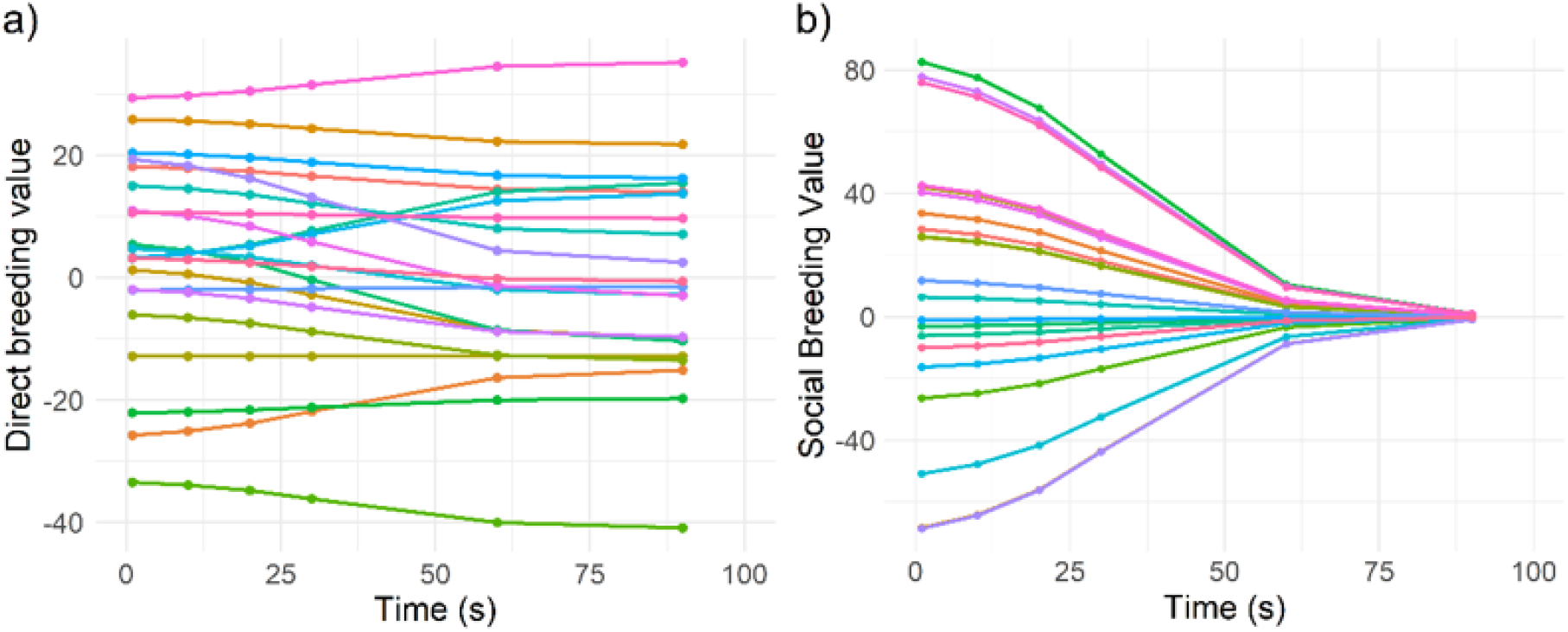
Progression of direct breeding values (a) and social breeding values (b) within animal. Each line represents a progression of a unique animal. Only 20 random animals were used to create the plot.

### Variance component conversion from visit approach to total phenotypes

Variance components of the total phenotype TFIP (the total feed consumption over the whole testing period) were approximated from variance components obtained from a visit-based approach using FIV (see Appendix). The reconverted variances are shown in Table 2. We used median values of the number of visits and median values of the covariate *Φ* to approximate the variance of a representative animal. When using Model 2, the resulting reconverted direct heritability was 0.25, and the social heritability (equal to the proportion of the phenotypic variance explained by the social variance) was 0.55. When model 3 was applied, direct heritability was 0.34 and social heritability was zero. The reconverted genetic correlation between direct and social effects was almost zero in both models 2 and 3.

**Table 2.** Comparison between reconverted variances from visit approach to total intake (RR Model) and variance components estimation of total consumption over the testing period.

| Component | Reconverted from<br>RR Model 2 | Reconverted from RR<br>Model 3 |
| --- | --- | --- |
| $\sigma_{pen}^2$ | 513 | 505 |
| $\sigma_{u_d}^2$ | 1342 | 1326 |
| $\sigma_{u_s}^2$ | 2882 | 9.52 |
| $\sigma_{u_d u_s}$ | -5.71 | 0.87 |
| $\sigma_{pe_d}^2$ | 3114 | 1937 |
| $\sigma_{pe_s}^2$ | - | 1655 |
| $\sigma_{pe_d pe_s}$ | - | -49.5 |
| $\sigma_e^2$ | 25.3 | 25.3 |
| $\sigma_{phe}^2$ | 5227 | 3804 |
| $h_d^2$ | 0.25 | 0.34 |
| $h_s^2$ | 0.55 | 0.00 |
| $rg(u_d u_s)$ | -0.00 | 0.03 |

## Discussion

### Random regression analysis

We modelized social genetic effects (model 2 and 3) using data from automatic feeders. Model 2 includes social genetic effects and assumes that both additive and social genetic effects are influenced by the interval time between visits (interval time between the entry of the follower and the exit of the pig occupying the feeder). Model 3 adds a permanent environmental effect for the follower. Also, a classical genetic model without social genetic effects (model 1) was also used as a reference. Random regression analyses applied in Models 2 and 3 allowed us to evaluate how direct additive, social additive, and permanent environmental effects change as a continuous function of the covariate *ϕ*, which represents the interval between feeder visits. This approach expands the one proposed by Angarita et al. (2021), which we had previously applied to this same Iberian pig population (Nuñez et al., 2025) using discrete time-interval classes rather than a continuous covariate. In that earlier study, differences were described between extreme time intervals categories: less than 60s and longer than 600s. The random regression approach used here overcomes this limitation by allowing the genetic and environmental effects to be modelled continuously across the entire range of the gradient, rather than being restricted to discrete classes without the need of using sub datasets. Although this covariate was constructed based on the social variance, applying it also to the direct additive component allows us to determine how susceptible an animal may be to the “social pressure”. As described above, the covariate *Φ* was constructed by fitting a function to the squared root of the normalized social variance, estimated from sub-datasets defined by different time intervals. This approach appears appropriate and coherent within this particular population; however, applying it to a different breed would require fitting a new function, since the relationship between social variance and time interval could differ across genetic backgrounds.

In Model 1, the pen variance was relatively large, suggesting that part of this variance is capturing social sources of variation. The random regression models (2 and 3) reduced the residual variance. Model 3, which additionally included a social permanent environmental effect, provided the best statistical fit according to the likelihood values. The magnitude of the combined social genetics and social permanent environment variances indicates that the model including the follower is an adequate model, given that it explains a large part of the variation, reduces residual variance, and is statistically significant.

However, the asymptotic correlation in Model 3 between the social genetic and social permanent environmental slope variances was high (−0.60), indicating that these two effects could not be reliably disentangled, likely due to the data structure. Consequently, although Model 3 achieved the best statistical fit, the biological interpretation of these two components is uncertain. Therefore, we focused primarily on Model 2. While the estimated social genetic variance in this model is expected to be inflated because it also captures part of the social permanent environmental variation, it indicates that a substantial proportion of the variation associated with the follower effect contributes to differences in feed intake per visit.

The results from variance component estimation denote that social effects are maximum when time interval is zero, but decrease quickly, being irrelevant when time interval exceeds 60s. This is as expected *a priori*. To the best of our knowledge, this visit-based approach is original and was only used in Angarita et al. (2021) and Nuñez et al. (2025). Both studies showed that the proportion of the phenotypic variance explained by social effects dropped to less than 1% in long-time intervals. Many previous studies used traditional social models (Bijma et al., 2007) where the phenotype was collapsed into a unique observation per animal and time period, namely daily feed intake, showing relevant increases in the proportion of the genetic variance when social indirect effects are included (Bergsma et al., 2008; Herrera-Cáceres et al., 2019; Kavlak et al., 2021; Nielsen et al., 2018). Both approaches (this work and the later preceding authors) support the importance of social effects on feed intake when animals are housed in groups.

No previous studies were found using random regression to fit direct and social effects as a function of the time interval. Thanks to this approach, we obtained direct-social additive genetic correlations at different time intervals. The weak correlation (very close to zero) indicates that simultaneous selection based on direct additive and social breeding values is feasible. However, careful consideration is needed, as this correlation may vary under different conditions. For example, under more competitive environments, such as feed restriction or high density, social genetic variance may become more relevant, and the direct–social covariance is expected to be more negative (Piles et al., 2017). Under these socially challenging scenarios, animals selected only for estimated direct breeding values 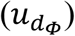 may become more competitive, which tends to reduce the mean phenotypic value of the pen.

### Direct and total correlations between timepoints

The results from correlations between timepoints indicate that 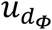 are highly stable across the gradient, with correlations ranging from 0.92 to 1. In contrast, total genetic correlations showed a marked decrease over time, dropping to 0.74 at 60 s (point from which the social genetic variance is negligible) and 0.62 at 120 s. Social genetic correlations were not reported because only a random regression slope was fitted for the social effect. As a result, each animal was characterized by a single social random coefficient, yielding a correlation of 1 across all time points.

### Intra-animal variation on estimated direct and social breeding values

The most important variation in 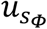 between animals was observed in short intervals, being reduced and staying constant after 60 s. This result is consistent with the estimation of social genetic variance. Based on the biology of social effects, it seems reasonable to focus on social genetic effects only when time intervals are short, as no biologically meaningful social effect is expected over long-time intervals between visits. Social effects estimated over long periods do not make biological sense and may introduce noise into the estimations. Our results suggest that a subset of data including only visits with time intervals between 0 and 30 s may be enough for estimating social effects, whereas longer intervals may lead to spurious results.

### Visit vs overall approaches

The visit-based approach applied in this study differs from the traditional approach proposed by Bijma et al., (2007), where a single aggregated phenotype of an animal (e.g., DFI) depends on its direct additive genetic effects and on the social genetic effects of its (*n* − 1) pen mates. In contrast, the visit-based approach models each visit individually, estimating 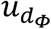 and 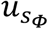 based on all recorded visits and accounting not only for different followers within the pen but also for their frequencies. An important advantage of the visit-based approach is its easier implementation, as the social effects of each observation depends only on the genetic effect of the follower (only one column on the data) being similar to a model with maternal effects (Willham, 1963; Cheverud, 1984). The visit-based approach is computationally easier and of faster convergence due to the increased precision provided by repeated measures (Jamrozik & Schaeffer, 1997). However, despite its advantages, the visit-based approach remains being a trait per visit, which is not of direct economic importance for farmers.

For the reason, we aimed to convert the variance components estimated for feed intake per visit (FIV) into a variance of total feed intake (TFIP). This approach allows us to benefit from the advantages of the visit-based model while still being able to evaluate how the overall phenotype is affected by each random effect. The results show that the relatively small proportion of additive variance estimated with the visit-based approach (~3.5%) translates into a substantially larger proportion of the variance in the aggregated phenotype (~25%). Similarly, the social component represented ~8% of the variance at the visit level and accounted for a larger proportion (55%) of the variance in the aggregated phenotype. These results highlight the potential importance of follower effects for overall feed intake. However, because the model does not include a social permanent environmental effect, the estimated social component cannot be unequivocally attributed to genetic effects and may include both genetic and environmental sources of social variation.

## Conclusion

In conclusion, this study provides a visit-based framework to identify and quantify social interactions between pigs according to the time interval between feeding events. By incorporating interaction strength into a random regression model, we show that social genetic effects are strongly dependent on the temporal structure of interactions, whereas additive genetic effects remain relatively stable. These results demonstrate that individual feeding events can provide information on the genetic basis of social interactions without assuming that all group members contribute equally to the social environment.

Furthermore, the proposed framework allows variance components estimated from individual feeding events to be aggregated into genetic and environmental variance components for total feed intake over the growth period. The contribution of social effects to the aggregated trait supports their relevance beyond individual feeding events and illustrates how detailed measurements of social interactions can be connected to a phenotype of practical interest.

Overall, this approach provides a more flexible framework for modelling social interactions. Although demonstrated here for feed intake in pigs, the framework could be extended to other continuously recorded traits and group-housed individuals where interactions vary in strength over time.

## Data availability statement

All the data are available at https://doi.org/10.6084/m9.figshare.33428779.

## Acknowledgments

The authors thank IngaFood S.A for the data.

## Appendix

### Derivation of total phenotypic variance from visit-based approach

In our random regression applied to visit approach for animal *i*, the direct genetic value was obtained as, 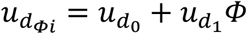, so the breeding value of the total feed intake can be written as 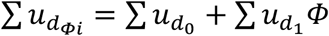, being the additive variance equal to 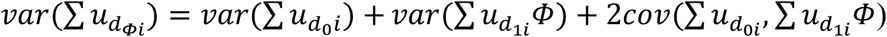. The previous equation can be separated into the three main terms:

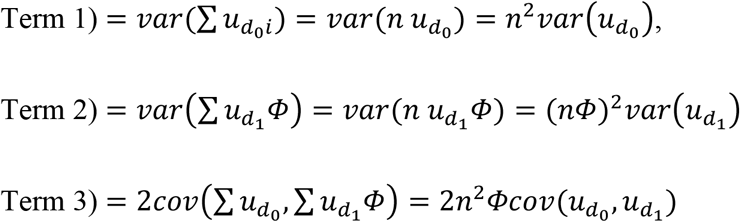

Now, the additive variance of the global feed intake can be approximated by:

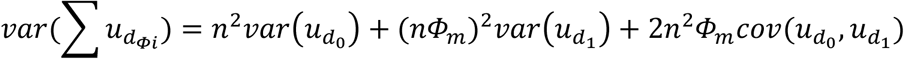

Assumptions:

- Breeding values for the same animal are constant, 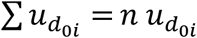.
- *n* is the median number of visits, and *Φ*_*m*_ the median value of the covariate *Φ* across the total visits. These assumptions aimed to approximate the reconstructed variance to a representative animal.

The social variance affecting individual *i* is approximated into the global social variance as follows:

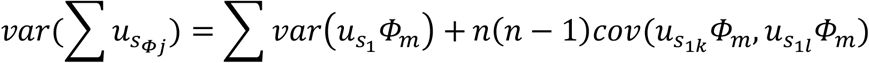

where *k* and *l* represent two different visits of individual *i*, so 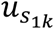 and 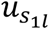 belong to two different *j* followers. Then,

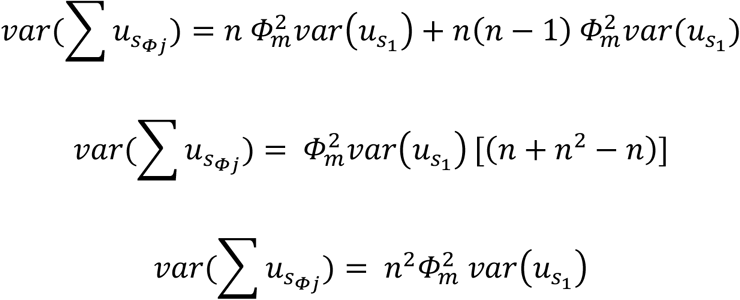

And for random permanent variance (model 2), where:

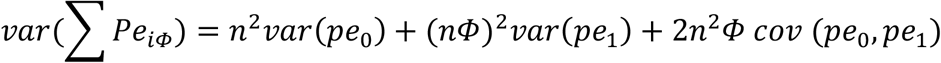

In contrast, the residual variance of the global phenotype is different because the residuals change between visits (∑ *e*_*i*_ ≠ *n e*_*i*_). Here, the variance of a sum does apply, where:

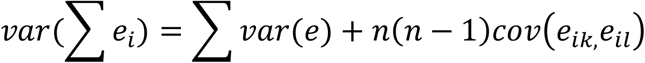

As the residuals from different visits are considered independent, *cov*(*e*_*ik*,_*e*_*il*_) = 0 and

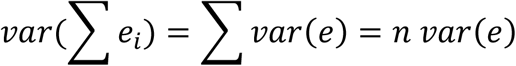

The phenotypic variance of the global phenotype was approximated as:

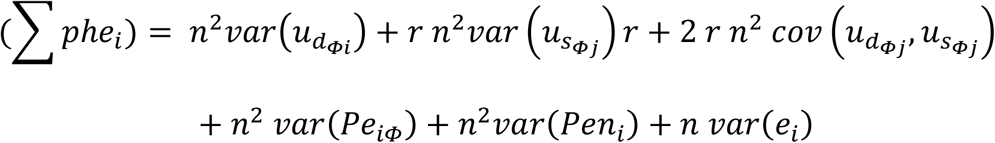

Note that, when the phenotypic variance of animal *i* is computed, *u*_*d*_ and *u*_*s*_ belong to different animals. The value of *u*_*d*_ for the phenotype of animal *i* is constant, while the value of *u*_*s*_ changes, as the follower *j* changes from one visit to another. For this reason, the reconstructed social variance and direct-additive covariance includes the kinship relationship (*r*) between the different followers.

Being the 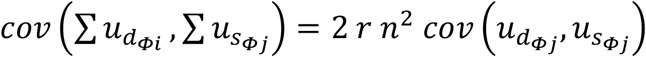 which can be decomposed into

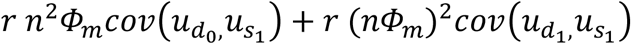

